# Early visual cortex tracks speech envelope in the absence of visual input

**DOI:** 10.1101/2022.06.28.497713

**Authors:** Evgenia Bednaya, Bojana Mirkovic, Martina Berto, Emiliano Ricciardi, Alice Martinelli, Alessandra Federici, Stefan Debener, Davide Bottari

## Abstract

Neural entrainment to continuous speech is typically observed within the language network and can be modulated by both low-level acoustic features and high-level meaningful linguistic units (e.g., phonemes, phrases, and sentences). Recent evidence showed that visual cortex may entrain to speech envelope, however its putative role in the hierarchy of speech processing remains unknown. We tested blindfolded participants who listened to semantically meaningful or meaningless stories, either in quiet or embedded in multi-talker babble noise. Entrainment to speech was assessed with forward linear modeling of participants’ EEG activity. We investigated (1) low-level acoustic effects by contrasting neural tracking of speech presented in quiet or noise and (2) high-level linguistic effects by contrasting neural tracking to meaningful or meaningless stories. Results showed that envelope tracking was enhanced and delayed for speech embedded in noise compared to quiet. When semantic information was missing, entrainment to speech envelope was fastened and reduced. Source modeling revealed that envelope tracking engaged wide neural networks beyond the auditory cortex, including early visual cortex. Surprisingly, while no clear influence of semantic content was found, the magnitude of visual cortex entrainment was affected by low-level features. The decrease of sound SNR-level dampened visual cortex tracking, suggesting an active suppressing mechanism in challenging listening conditions. Altogether, these findings provide further evidence of a functional role of early visual cortex in the entrainment to continuous speech.

## Introduction

Neuronal populations developed the ability to synchronize their activity (through aligning the phase) to temporal regularities of a continuous input (Lakatos et al., 2019; Obleser & Kayser, 2019). This neural entrainment influences several aspects of processing, including language. In this context, neural activity entrained to amplitude modulations over time of continuous speech (that is, the envelope) has been consistently reported (Ding & Simon, 2014). The exact functional meaning of the entrainment to the speech envelope is still unclear. Several studies showed that intelligible speech is not mandatory for neural tracking (Howard & Poeppel, 2010; Luo & Poeppel, 2007). However, during comprehension, phase-locked responses to speech in the auditory cortex are enhanced (Gross et al., 2013; Peelle et al., 2013). Moreover, entrainment to an attended speaker’s speech envelope in noisy environments appears to play a role in solving the so-called “cocktail party” (Cherry, 1953) problem (Ding, Chatterjee, & Simon, 2014; Riecke, Formisano, Sorger, Başkent, & Gaudrain, 2018). Based on this evidence, entrainment to speech envelope may be involved in promoting the perception of linguistic information (Poeppel & Assaneo, 2020) and facilitating speech comprehension (Ahissar et al., 2001; Luo & Poeppel, 2007), especially in challenging acoustic environments (e.g., Kerlin, Shahin, & Miller, 2010; Zion Golumbic et al., 2013). Importantly, neural entrainment to temporal dynamics of speech is modulated by low-level acoustic features (Ding et al., 2014) and high-level meaningful linguistic units, such as phonetic information, phrases, and sentences (Di Liberto, O’Sullivan, & Lalor, 2015).

Neural entrainment does not only occur for the auditory input of speech (A. E. O’Sullivan, Crosse, Liberto, Cheveigné, & Lalor, 2021; Plass, Brang, Suzuki, & Grabowecky, 2020). Recent magnetoencephalography (MEG) studies revealed that the early visual areas entrain even to silent lip movements (Bourguignon, Baart, Kapnoula, & Molinaro, 2018, 2020; Hauswald, Lithari, Collignon, Leonardelli, & Weisz, 2018). This neural tracking is modulated by audiovisual congruences and boosts speech comprehension in noisy conditions (Park, Kayser, Thut, & Gross, 2016). The contribution of visual cortices in language processing is not limited to visual or audiovisual representations of spoken language. There is scattered evidence that the early visual cortex is also active during purely auditory stimulation (Brang et al., 2022; Petro, Paton, & Muckli, 2017; Vetter, Smith, & Muckli, 2014) and while listening to spoken language (e.g., Martinelli et al., 2020; Seydell-Greenwald, Wang, Newport, Bi, & Striem-Amit, 2021; Wolmetz, Poeppel, & Rapp, 2011). Importantly, such activations cannot be explained by semantic-based imagery alone but rather seem to reflect genuine responses to language input; in fact, the visual cortex also responds to abstract concepts with low imaginability rates (Seydell-Greenwald et al., 2021). Overall, this evidence highlights a putative role of the visual cortex in mapping temporal modulations of incoming sounds, especially in the absence of competing retinal input (Martinelli et al., 2020; Vetter et al., 2014). However, the exact role of the visual cortex in the hierarchy of speech processing remains unclear.

Here, we investigated the neural tracking of speech envelope when visual input is absent. Using electroencephalography (EEG), we recorded neural responses of blindfolded individuals while they were listening to stories presented in isolation (Quiet) or combined with multi-talker babble noise at different signal-to-noise ratios (SNR; Noise). Stories comprised either meaningful (speech) or meaningless (jabberwocky) narration. We used a temporal response function (TRF) to model neural tracking of broadband speech envelope (in 2-8 Hz range; as in: Hausfeld, Riecke, Valente, & Formisano, 2018; Mirkovic, Debener, Jaeger, & De Vos, 2015; J. A. O’Sullivan et al., 2015). TRF approach allows linear mapping between neurophysiological responses and continuous speech stimuli (Crosse, Di Liberto, Bednar, & Lalor, 2016; Crosse et al., 2021) and has been used to measure entrainment to speech in both clear and challenging listening conditions (e.g., Decruy, Vanthornhout, & Francart, 2019; Di Liberto et al., 2015; Ding et al., 2014; Ding, Melloni, Zhang, Tian, & Poeppel, 2016; Ding & Simon, 2014; Legendre, Andrillon, Koroma, & Kouider, 2019; J. A.O’Sullivan et al., 2015).

To disambiguate the effects of lower-level acoustic and higher-level linguistic processing using continuous naturalistic stimuli, we built a hierarchical model. We specifically assessed the effects of (i) low-level acoustic features by contrasting TRFs resulting from listening to stories presented in quiet vs. in noise, and (ii) high-level linguistic information by contrasting TRFs resulting from listening to meaningful (speech) vs. meaningless (jabberwocky) stories, both embedded in noise. Finally, we tested how low-level and high-level information effects are distributed at the source level, with a focus on whether and how speech envelope information is mapped in the visual cortex in the absence of competing visual information.

## Materials and Methods

### Participants

Nineteen native speakers of the Italian language took part in the study (N = 19; age: median = 28; min = 22; max = 32; females = 12; all right-handed). We excluded one participant because of an error in the presentation script during EEG acquisition and three more participants due to their inability to complete the experiment, resulting in a final sample of fifteen participants (N = 15; age: median = 28; min = 22; max = 30; females= 10). All participants self-reported the absence of any hearing problems and neurological disorders. The experimental protocol was approved by the local ethics committee and conducted following the Declaration of Helsinki. All participants were informed in advance that they would be blindfolded during the experiment, signed written informed consent prior to the study, and received monetary compensation for their participation.

### Stimuli

We used two types of target stories: (i) meaningful (speech) and (ii) meaningless (jabberwocky) narration. Meaningful stories were extracted from the fiction book for teens *Polissena del Porcello* by (Pitzorno, 1993). Meaningless stories were extracted from the books containing nonsense, metasemantic (jabberwocky) poems and texts: *Gnòsi delle fànfole* by (Maraini, 2019) and *Esercizi di Stile* by (Queneau, 1947/1983). Note that syntactic information is preserved in jabberwocky stories, whereas semantic information is absent or significantly reduced.

Target stories were narrated by a trained Italian actress. We registered stories in a soundproof booth, using a video camera with an external condenser microphone (Olympus ME51S) at sampling frequency of 48000 kHz. To create stimuli for our EEG experiment, we extracted the audio material from the recorded files and edited them in Audacity® software (version 2.3.0, https://www.audacityteam.org/) and with a custom code using Signal Processing toolbox incorporated in MATLAB (version R2018b, Natick, Massachusetts: The MathWorks Inc.). Specifically, we: (i) inspected raw audio files for pronunciation errors and long breaths, consequently removing them, (ii) downsampled audio to 44100 Hz, set to 16-bit and converted from Stereo to Mono, (iii) truncated long pauses and silent periods exceeding 0.5 s to 0.5 s, (iv) trimmed resulting files to the same length (∼ 15 min), (v) identified the noise floor of the frequencies comprising the noise via “Get Noise Profile” feature and subsequently removed low-amplitude background noise using the Noise Reduction built-in feature based on an algorithm using Fourier analysis, (vi) normalized resulting files to the same common root-mean-square (RMS) value to ensure no variation of loudness across stories. Natural variations of loudness within each story were preserved.

We combined the target stories with a five-talker babble to construct stimuli in which the target story was embedded in the noise. Here, we used the babble noise, which is a non-stationary noise that works well both as an energetic and informational masker, efficiently reducing intelligibility and speech quality (Brungart, 2001; X. Wang & Xu, 2021). The babble noise was a mixture of five different voices (2 females, 3 males, all native Italian speakers). Every speaker was recorded in the soundproof booth, reading several non-related extracts from the fiction book *La Strada* by (McCarthy, 2006/2014). These individual recordings were registered and edited with the similar routine described above for the target stimuli. Then, individual recordings were superimposed, resulting in multi-talker babble. Finally, the initial 500 ms of the multi-talker babble got discarded to eliminate a part that did not contain all five talkers.

The first 5 s of the resulting multi-talker babble were set to zero/”muted,” followed by 5 s of fade-in to make it easier for the participants to identify and track the target stories in the multi-talker babble noise. Then, with custom MATLAB scripts, we normalized the target stories and the babble to a common RMS value to make sure there would be no story or any of its segments standing out from the noise, and then superimposed the stories and the babble at two SNR levels (SNR1 = +3.52 dB, – for both meaningful and meaningless stories, and SNR2 = +1.74 dB, – for meaningful story only; see supplementary material for more details). As the last step, we normalized all the resulting audio files for all conditions once again to a common RMS value to achieve equal loudness across the stimuli and consequently verified each file’s spectrogram in Audacity.

Altogether, we constructed stimuli to generate four experimental conditions: 1) *Speech-in-Quiet*, 2) *Speech-in-Noise at SNR1*, 3) *Speech-in-Noise at SNR2*, and 4) *Jabberwocky-in-Noise at SNR1* (Figure 1A). Each experimental condition contained a particular story divided into three parts of ∼ 5 min, therefore the total duration of continuous speech stimuli per condition was ∼ 15 min.

**Figure 1.**
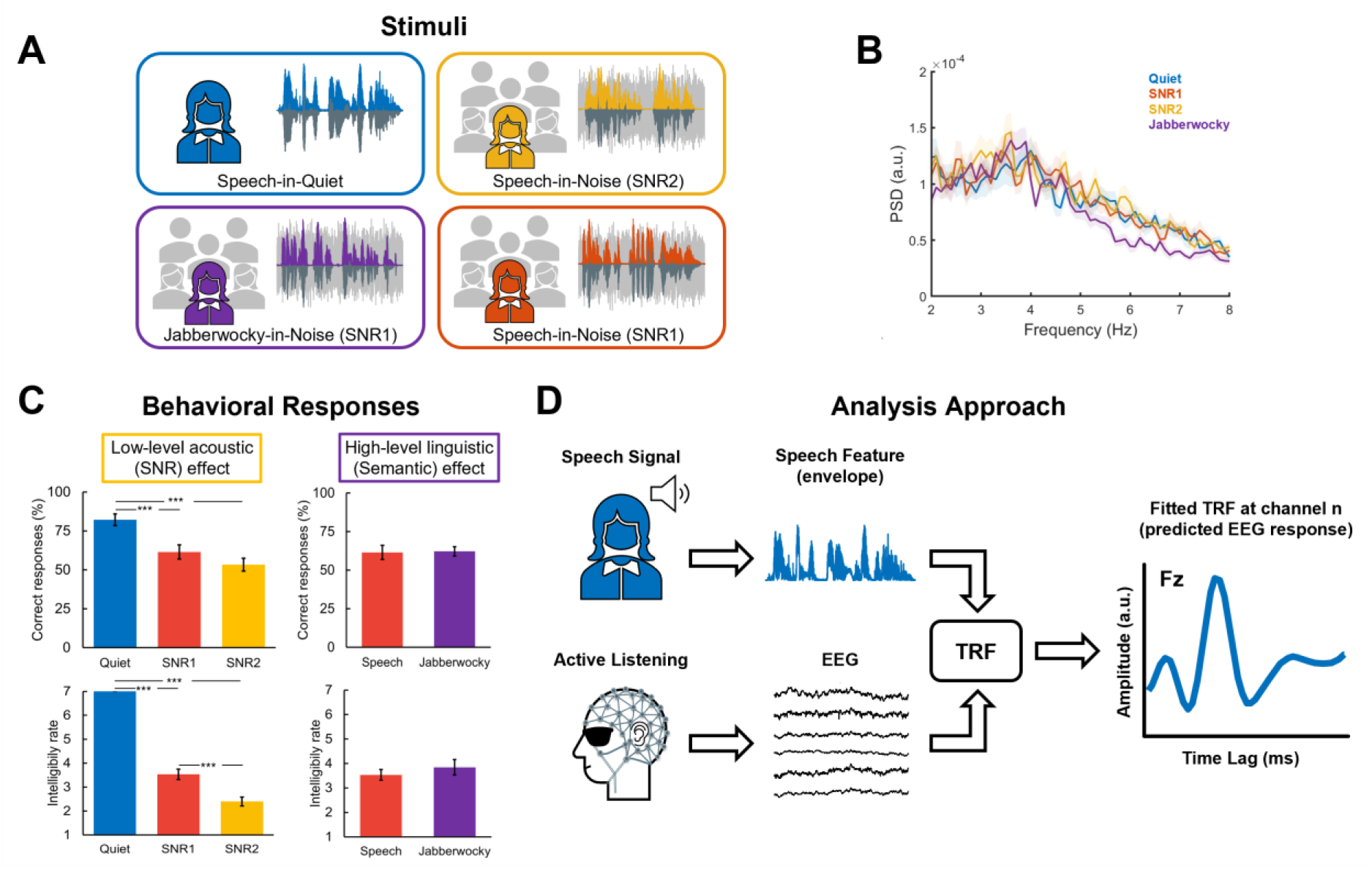
Stimuli, Behavioral Responses, and Analysis Approach. **(A)** Stimuli consisted of continuous (i) meaningful (Speech) and (ii) meaningless (Jabberwocky) stories presented either in quiet (Quiet) or as embedded in the multi-talker babble noise at a different signal-to-noise ratio (SNR1; SNR2). The babble noise was a mixture of five voices (2 females, 3 males) reading extracts from a book. The acoustic envelopes were extracted for further analysis through the Hilbert transform and filtering in the range between 2 and 8 Hz. **(B)** Power spectra density estimates of normalized acoustic envelopes were obtained using Welch’s method with a 10 s Hamming window and half-overlap. Bold lines indicate average across trials; shaded areas indicate standard error of the mean. **(C)** Behavioral Responses represented by correct responses (Top) and intelligibility rates (Bottom). Barplots display mean ± SE across participants. Asterisks indicate statistically significant differences (***p < 0.001). **(D)** Neural tracking of the speech envelope was estimated using the forward encoding approach – Temporal Response Function (TRF). Ridge regression-based linear models (TRFs) were fitted to participants’ neural data, obtained during active listening, to predict EEG response for of a given EEG channel from speech envelope.

To test the effect of low-level acoustic (SNR) information, we compared neural tracking in *Speech-in-Quiet* condition and *Speech-in-Noise at SNR2* condition. To test the effect of high-level linguistic (semantic) information, we compared neural tracking in *Speech-in-Noise at SNR1* condition and *Jabberwocky-in-Noise at SNR1* condition.

### Task and Experimental Procedure

Participants performed four blocks, each consisted of one experimental condition. During the first block, they always listened to the story without background noise (i.e., *Speech-in-Quiet* condition). This was done to help the participants habituate both to the (target) narrator’s voice and the experimental design since this condition was the easiest to attend. The order of the remaining three blocks was randomized across participants. Each of the four blocks consisted of a story that lasted ∼ 15 min divided into three parts ∼ 5 min (see supplementary material for further details). Participants listened to each part of the story only once, without repetition, therefore avoiding the possibility of predicting the content of the story. To maintain the continuity of the storyline within each block, each part within each story followed the previous part chronologically.

We instructed participants to attentively listen to the target story (narrated by the female voice and guided by the first 5 s of the audio) while ignoring babble noise in the background. To ensure that the participants were actively attending to the stimuli, at the end of each part, they answered three specific Yes/No questions about the part of the story that they just listened to (for example, “*Il cane di Lucrezia è un San Bernardo?* [Is Lucrezia’s dog a Saint Bernard?]”; see supplementary material for the full list of questions). If they were not sure about the correct answer between the two, they had to choose the answer that seemed to them the most probable. To answer, participants pressed corresponding buttons on the response panel with their index and middle fingers.

At the end of each part, we asked participants to self-report intelligibility rates of the target story on a Likert scale (where 1 – absolutely non-intelligible, 7 - very intelligible) and let them have a short break lasting ∼ 2 min. We also ensured that none of the participants was familiar with or recently exposed to the target stories. Moreover, we informally assessed a participant’s comfort, alertness, and motivation to continue the experiment during short and long breaks. We removed the blindfolding mask during the breaks between each blocks (every 15 minutes) for the participants’ comfort and in order to avoid inducing short term cross-modal plasticity effects resulting from the prolonged visual deprivation (Landry, Shiller, & Champoux, 2013; Lazzouni, Voss, & Lepore, 2012; Merabet et al., 2008).

The experiment was controlled with E-Prime® software (version 3.0, Schneider et al., 2002). All instructions and speech stimuli were presented through a single front-facing loudspeaker (Bose Companion® series III multimedia speaker system, USA) placed in front of the participants at approximately 1 m distance from their heads. Stimuli were delivered at ∼ 60 dB sound pressure level (SPL), measured at the participant’s ear, and reported by all the participants as comfortable volume.

To accurately measure the actual onset time of our stimuli, we administered a timing-test using Audio/Visual (AV) Device (Electrical Geodesics, Inc.) compatible with E-Prime software and NetStation system. The measured average delay in time was constant and about + 5 ms regarding the stimulus onset.

### EEG Recording

Before starting the experiment, each participant received a brief instruction and had a short (∼ 1 min) “training” session on how to control over muscular artifacts through monitoring their EEG signal displayed on the computer screen. Then, we applied the blindfolding mask to the participant, and they were reminded to keep their eyes open during the EEG recordings, though blinking was permitted whenever they wanted. Moreover, we recorded resting-state EEG data for about 2 minutes at the beginning of each experiment while the participant kept their eyes open. Obtained resting-state data served as calibration data to attenuate EEG artifacts during the preprocessing step.

During the tasks, the participants were seated comfortably in a chair in a dark, soundproofed booth (BOXY, B-Beng s.r.l., Italy). The EEG recordings were acquired at a sampling rate of 500 Hz using NetStation5 software together with a Net Amps 400 EGI amplifier connected to 64 electrodes HydroCel Geodesic Sensor Net (Electrical Geodesics, Inc.), all signals referenced to vertex (additional channel E65/Cz). For data visualization purposes only, the data were band-pass filtered online using the digital filter from 1.0 to 100 Hz, and online digital anti-alias filter aligning EEG recordings with real-time events was kept on. Electrode impedances were kept below 50 kΩ and were checked between the blocks (when the blindfolding mask was reapplied).

Participants were encouraged to take a break after each block and get enough rest before continuing. They also were reminded about the importance of staying attentive, keeping eyes open while blindfolded, and avoiding excessive movements during the EEG recordings.

### EEG Preprocessing

We preprocessed continuous EEG raw data offline using custom MATLAB (version R2018b, Mathworks Inc., Natick, MA) scripts together with EEGLAB toolbox (version 14.1.2b, Delorme & Makeig, 2004) for MATLAB.

First, the EEG data were submitted to cleaning with Artifact Subspace Reconstruction (ASR) - an automated artifact attenuation algorithm (clean_rawdata plug-in, version 2.1) for EEGLAB toolbox. We applied the default flatline criterion of 5 s, together with default transition band parameters ([0.25 0.75]). ASR algorithm was chosen due to its objective and reproducible evaluation of artifactual components in EEG data. ASR is based on Principal Component Analysis (PCA) sliding window and effectively attenuates high-variance signal components in the EEG data (including eye blinks, eye movements, and motion artifacts). Specifically, first, the algorithm automatically identifies the most artifact-free part of the data (here, the resting-state data) to use it as the calibration data to compute the statistics. Next, a 500 ms PCA sliding window with 50% overlap is applied across all the channels to identify “bad” principal components. Then, the algorithm identifies the subspaces in which the signal exceeds 5 standard deviations away from the calibration data as corrupted and rejects them. Finally, it reconstructs the high variance subspaces using a mixing matrix calculated based on the calibration data.

The artifact attenuated EEG data were preprocessed as follows: (i) re-referenced from E65/Cz electrode to a common average reference, (ii) band-pass filtered from 0.1 to 40 Hz (low-pass: FIR filter, filter order: 100, window type: Hann; high-pass: FIR filter, filter order: 500, window type: Hann), (iii) downsampled to 250 Hz, (iv) epoched according to the onset of acoustic stimuli (related to each part of the story), adjusting to measured +5 ms onset delay in time and discarding the first 5 s of target-speech alone and 5 s of fade-in for the babble noise, (v) band-pass filtered between 2 and 8 Hz (filter type and parameters the same as described above), (vi) downsampled to 64 Hz, (vii) EEG data corresponding to each of the three ∼ 5 min parts of the story were concatenated, (viii) and segmented into 1 min long trials, resulting in 12 trials per block per subject (N = 12). The preprocessed EEG data for each trial were z-scored to optimize cross-validation procedure during encoding (Crosse et al., 2016).

### Extraction of Acoustic Envelope

First, audio files containing relevant parts of the target stories were concatenated and segmented into corresponding 1 min long trials, resulting in 12 trials per speech envelope per subject (N = 12) (Figure 1B). Next, the acoustic envelope per each trial was obtained taking the absolute value of the Hilbert transform of the original target stories (i.e., without babble noise) followed by a low-pass filtering using a 3rd-order Butterworth filter with a cut-off frequency of 8 Hz (filtfilt function in MATLAB) and downsampling the resulting signal to 64 Hz, so to be matched with the EEG data (e.g., Mirkovic et al., 2015; J. A. O’Sullivan et al., 2015). Finally, the resultant extracted envelopes were normalized by dividing by maximum value.

### Estimation of TRF

We modeled where and how the neural response to the speech envelope of the target stories is encoded in the brain, using a linear prediction approach known as temporal response function (TRF) (Figure 1D). The TRF approach, incorporated in mTRF toolbox (Crosse et al., 2016), allows to predict previously unseen EEG response from the stimulus and has been used to model the neural tracking of acoustic and linguistic properties of naturalistic continuous speech (Drennan & Lalor, 2019; Obleser & Kayser, 2019).

The TRF is a mathematical function that is based on the ridge regression and could be described as follows:

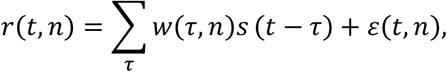

where *t* = 0, 1, … T is time, *r*(*t, n*) is the EEG response from an individual channel, *s*(*t*) is the stimulus feature(s) (e.g., speech envelope), *τ* is the range of time-lags between *s* and *r, w*(*τ, n*) are the regression weights over time-lags, and *ε*(*t*) is a residual response at each channel not explained by the TRF model (Crosse et al., 2016). Specifically, TRF can be viewed as a filter that describes the linear relationship between a continuous speech stimulus and a continuous neural response for a specified range of time-lags related to stimulus occurrence (Crosse et al., 2016).

The important assumptions about the TRF include the fact that it reflects the same neural generators as cortical auditory evoked potentials (CAEPs) resulting in their comparable topographies and that it can be used to measure neural tracking of speech envelope (Lalor & Foxe, 2010; Lalor, Power, Reilly, & Foxe, 2009). We fitted separate models (TRFs) to predict response in each EEG channel, using time-lags from -100 to 600 ms related to stimulus onset, typically used to capture CAEP components. Here we estimated the TRF using the envelope estimated between 2 and 8 Hz as previously performed (Legendre et al., 2019; Mirkovic et al., 2015; J. A. O’Sullivan et al., 2015).

The TRF models were trained using a leave-one-out cross-validation procedure, keeping all but one trial for training the model to predict EEG response from the stimuli and using a left-out trial for testing. Thus, a prediction model was obtained for every single trial, and then the final averaging across trials, within participants and conditions was performed, resulting in a grand average TRF model.

### Regularization Parameter Estimation

Regression models are exposed to overfitting the training data, that is, fitting the random noise rather than true relationships between variables and failing to generalize to unseen data. The problem of overfitting needs to be accounted for before making any interpretations from the resulting model since it could be misleading. Ridge regularization prevents the model from overfitting by penalizing the model weights, forcing them to be smaller, towards 0, so the model could become better generalized.

To control for model overfitting, we empirically identified the optimal regularization parameter (λ) of TRF models through leave-one-out cross-validation procedure, using a grid of ridge values (λ = {10^−6, 10^−5,…, 1, 10, …, 10^5, 10^6}), for time-lags from -100 to 600 ms. The regularization parameter λ was determined based on the mean squared error (MSE) value between the actual and predicted EEG responses. The optimal regularization parameter was the one yielding the lowest MSE on the testing data (here, identified as λ=10^3) and kept constant across channels, participants, and conditions allowing to generalize across them at the group level.

### Spatiotemporal Characteristics

Forward model weights are directly physiologically interpretable (Haufe et al., 2014) and allow us to get an insight about which channels contribute most to neural tracking of the speech envelope. The resulting topographical plots with TRF weights obtained per each individual time-lag window can be interpreted similarly to CAEPs in terms of both amplitude and direction (Lalor, Pearlmutter, Reilly, McDarby, & Foxe, 2006; Lalor et al., 2009). We investigated spatiotemporal characteristics of forward model weights by fitting the TRFs at different individual time-lags between the EEG response and the speech envelopes, using a sliding time-lag window of 45 with 30 ms overlap in a time-lag range from −115 to 620 ms. Finally, the estimate of forward model weights allowed us to directly transfer the data into source space avoiding further transformations (Haufe et al., 2014).

### Chance-level Estimation by Permutation Testing (Control)

To assess the ability of TRF models to predict neural responses (i.e., neural tracking) and verify that neural tracking was well above chance, we computed “null distributed” TRF model (Combrisson & Jerbi, 2015). We used a permutation-based approach with *mTRFpermute* function, incorporated in mTRF-toolbox (Crosse et al., 2016, 2021). Specifically, this approach cross-validates models, iteratively (1000 iterations) fitting TRFs on randomly mismatched pairings of speech envelopes/EEG responses and evaluating the models on matched data. This procedure was done separately for each trial, participant, and condition, and then grand averaged to get the average “null” TRF model, which served as a baseline (“control”).

### Source Estimation

Forward modeling allowed us to investigate the TRFs and better understand how the information about the envelope of continuous stimuli is encoded in the brain. Specifically, we tested how low-level (SNR) acoustic and high-level linguistic (Semantic) effects are distributed at sensor and source levels. Furthermore, we investigated whether and how the visual cortex is activated for neural tracking of the speech envelope in blindfolded individuals when competing retinal input is absent.

We performed source localization in Brainstorm software (Tadel, Baillet, Mosher, Pantazis, & Leahy, 2011) together with custom MATLAB scripts and the pipeline for EEG source estimation introduced by Stropahl and colleagues (2018; see also Bottari et al., 2020) that we adapted to the TRF data. Specifically, source localization was performed using dynamical Statistical Parametric Mapping (dSPM, Dale et al., 2000). A Boundary Element Model (BEM) was computed for each participant using default parameters to calculate the forward solution and constrain source locations to the cortical surface. We used a standard electrode layout together with a standard anatomy template (ICBM152) for all participants. The model resulted in a single dipole oriented perpendicularly to the cortical surface for each vertex since dipole orientations were constrained to the cortical surface. We did not perform individual noise modeling since TRF has no clear nor true baseline period. Instead, we used an identity matrix as a noise covariance matrix, with the assumption of equal unit variance of noise on every sensor.

We created visual regions of interest (ROIs) based on predefined scouts from the Destrieux atlas (Destrieux, Fischl, Dale, & Halgren, 2010) implemented in FreeSurfer (Fischl, 2012) and available in Brainstorm. Visual ROIs were selected for the left and right hemispheres and included primary (V1; Calcarine sulcus) and secondary (V2, Lingual gyrus) visual cortex, defined as the *‘S_calcarine’* and the *‘G_oc-tem_med-Lingual’* scouts in the atlas, correspondingly. These visual ROIs were selected based on recently reported evidence of their involvement in speech processing not only in blind but also sighted individuals, albeit to a lower extent (Martinelli et al., 2020; Petro et al., 2017; Seydell-Greenwald et al., 2021; Van Ackeren, Barbero, Mattioni, Bottini, & Collignon, 2018; Vetter et al., 2020, 2014). Upon the ROIs creation, their time-series were extracted and submitted to the analysis.

### Statistical Analysis

Participants’ behavioral responses concerning comprehension of the story were computed as the average correct responses (in %) across all three parts of the story, resulting in nine scores per participant for each condition. Intelligibility rates from each participant were computed similarly, by averaging across all three parts of the story. Statistical analysis of behavioral responses to assess low-level acoustic (SNR) effect was conducted using one-way repeated measure ANOVA. Post-hoc comparisons were made with two-tailed paired t-tests. Statistical analysis of behavioral responses to assess high-level linguistic (semantic) effect was performed using two-tailed paired t-tests.

As a sanity check, we first performed comparisons between the TRFs of each condition with the “null” TRF through paired t-tests, with the significance threshold set at p < 0.05 (one-tailed) and corrected for multiple comparisons with the false-discovery rate (FDR) at 0.05 (Benjamini & Hochberg, 1995), at two electrodes selected a priori on the midline frontocentral (Fz) and the occipital (Oz) scalp locations, over a range of post-stimulus time-lags between 0 and 600 ms

To access differences in the spatiotemporal profile of averaged TRFs between conditions, we performed non-parametric cluster-based permutation tests (Maris & Oostenveld, 2007) in FieldTrip toolbox (Oostenveld, Fries, Maris, & Schoffelen, 2011). A cluster was defined along *electrodes x time-lags* dimensions, with extension criteria set to at least two neighboring electrodes. The t-statistic for adjusted *electrode x time-lag* pairs exceeding a preset critical threshold of 5% (cluster alpha = 0.05) was summed, and the adjusted pairs formed the clusters. Then, two-tailed tests were performed at the whole brain level (across all electrodes and time-lags from 0 to 600 ms), using the Monte-Carlo method with 1000 permutations. The maximum of the summed t-statistic in the observed data was compared with a random partition formed by permuting the experimental condition labels, resulting in a critical p-value for each cluster. In case the cluster-based p-value was less than 0.025 (corresponding to a critical alpha level of 0.05 for two-tailed testing, accounting for both positive and negative clusters), we rejected our null hypothesis that there were no differences between TRFs for two conditions.

Finally, cluster-based statistics on sources at the whole-brain level were performed in Brainstorm, across all electrodes and time-lags from 0 to 600 ms, using Monte-Carlo method with 1000 permutations, alpha = 0.05, two-tailed (meaning alpha = 0.025 per each tail), cluster alpha = 0.05, and neighboring criteria for electrodes set for 2. Analysis of visual ROIs time-series between conditions was performed using paired t-tests, with the significance threshold set at p < 0.05 (one-tailed) and correcting for multiple comparisons with the FDR-method at 0.05.

## Results

### Behavioral responses

To ensure that the participants successfully understood the content of the target stories, they were asked to answer three Yes/No questions at the end of each segment (5 minutes). Moreover, participants were asked to self-rate the intelligibility of each part of the target story from 1 (absolutely non-intelligible) to 7 (very intelligible).

#### Low-level acoustic (SNR) effect

As expected, both comprehension scores and intelligibility rates gradually decreased with SNR (Figure 1C). Comprehension scores, converted to percentage of correct responses, decreased as a function of noise (*Speech-in-Quiet* mean ± SE: 82.22 ± 3.72%; *Speech-in-Noise at SNR1* mean ± SE: 61.48 ± 4.58%; *Speech-in-Noise at SNR2* mean ± SE: 53.33 ± 4.09%). A repeated measures ANOVA with a correction confirmed that listening condition significantly affected participants’ comprehension (F(2, 28) = 16.14, p = 0.00002, Huynh-Feldt corrected). Post-hoc comparisons showed that correct responses for *Speech-in-Quiet* were significantly higher than for *Speech-in-Noise at SNR1* (t(14) = 4.40, p = 0.0006) and *Speech-in-Noise at SNR2* (t(14) = 5.46, p = 0.0001), but no significant difference emerged between *Speech-in-Noise at SNR1* and *Speech-in-Noise at SNR2* (t(14) = 1.43, p = 0.17).

Intelligibility rates were in line with comprehension scores, dramatically dropping from *Speech-in-Quiet* (rated 7 by all participants, and thus reaching a ceiling which prevented comparisons with other conditions; see Liu & Wang, 2021; Šimkovic & Träuble, 2019) to *Speech-in-Noise at SNR1* (mean ± SE: 3.53 ± 0.22) and further significantly dropping at *Speech-in-Noise at SNR2* (mean ± SE: 2.40 ± 0.19; *Speech-in-Noise at SNR1* vs. *Speech-in-Noise at SNR2:* t(14) = 5.90, p < 0.0001).

#### High-level linguistic (Semantic) effect

We found no difference in correct responses and intelligibility rates between *Speech-in-Noise at SNR1* and *Jabberwocky-in-Noise at SNR1* (all p-values > 0.05; correct responses, mean ± SE: *Speech-in-Noise at SNR1:* 61.48 ± 4.58%; *Jabberwocky-in-Noise at SNR1:* 62.22 ± 3.03%; intelligibility rates, mean ± SE: *Speech-in-Noise at SNR1:* 3.53 ± 0.22; *Jabberwocky-in-Noise at SNR1*: 3.84 ± 0.32. Results indicated that participants were able to equally attend target stories embedded in noise (SNR1), regardless of semantic information.

### Neural tracking

#### Low-level acoustic (SNR) effect at the sensor level

First, we examined the temporal profile of SNR effect at preselected representative electrodes: frontal (Fz) and occipital (Oz; Figure 2A). The TRFs for *Speech-in-Quiet* and *Speech-in-Noise at SNR2* were significantly different from the “null” TRF (p < 0.05, one-tailed, FDR-corrected), suggesting that TRFs indeed reflected neural tracking of the speech envelope (Supplementary Figure S1).

**Figure 2.**
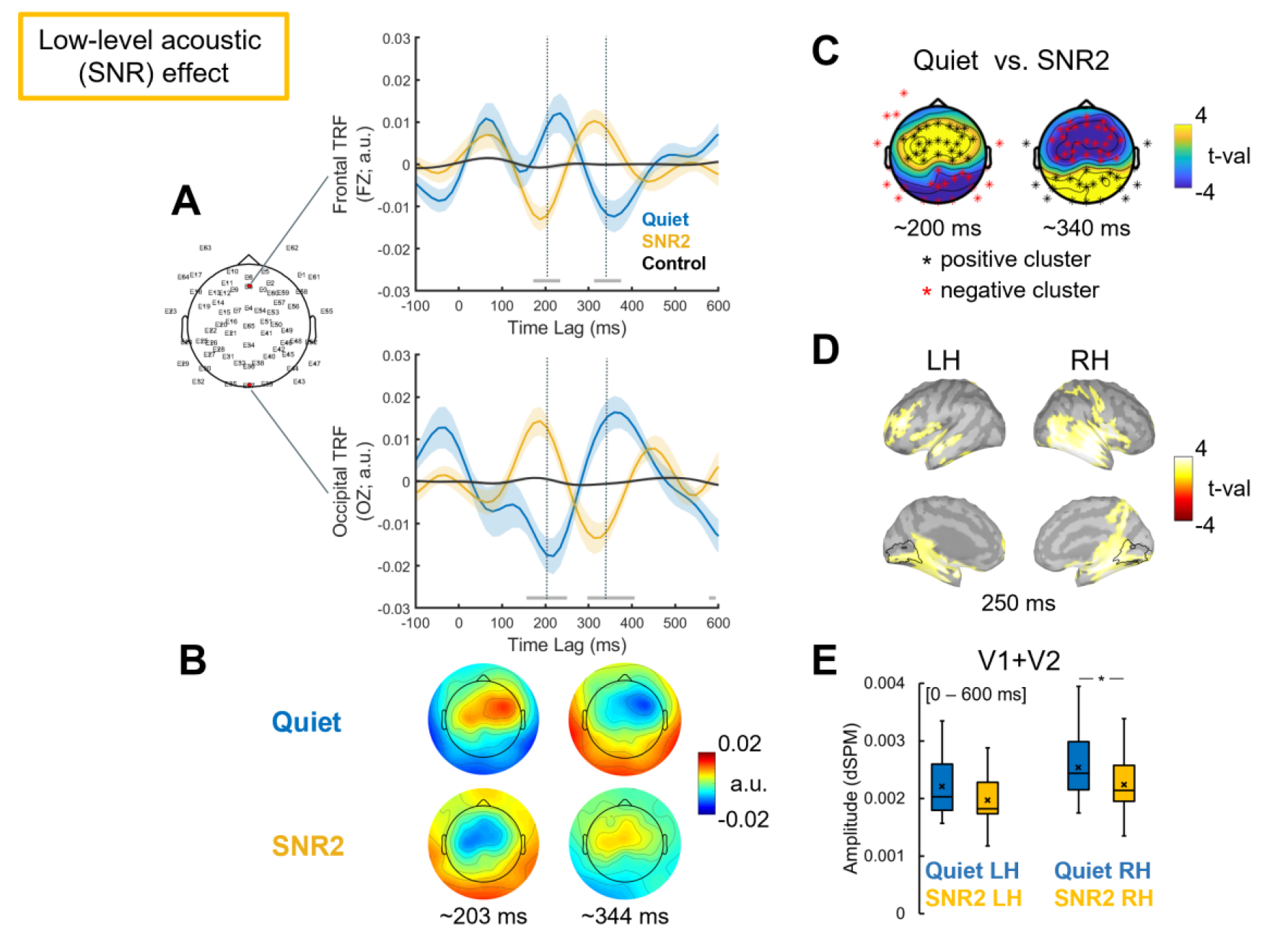
Low-level acoustic (SNR) effect. **(A)** Grand averaged temporal response functions (TRFs) for *Speech-in-Quiet* (Quiet, blue), *Speech-in-Noise at SNR2* (SNR2, yellow), and “null” TRF (Control, black). TRFs displayed over time-lags at frontal Fz and occipital Oz electrodes, marked with red on the electrode layout. Shaded areas represent the standard error of the mean (SE) across participants. Grey horizontal bars above the x-axis indicate time-lags at which TRFs for *Speech-in-Quiet* and *Speech-in-Noise at SNR2* differed significantly at these representative electrodes (series of paired two-tailed t-tests, p < 0.05, FDR-corrected). Grey dotted vertical lines indicate time-lags with the maximal difference between TRFs for *Speech-in-Quiet* and *Speech-in-Noise at SNR2*. **(B)** Topographic representations of TRFs, displayed at time-lags marked by grey dotted vertical lines on **A. (C)** The results of the cluster-based permutation test contrasting TRFs for *Speech-in-Quiet* vs. *Speech-in-Noise at SNR2*, displayed around time-lags marked by grey dotted vertical lines on **A**. Significant (p < 0.05, corrected for two tails, p < 0.025 for each tail) positive and negative clusters comprised the electrodes marked in black and in red asterisks, respectively. **(D)** Differences at the source level, contrasting TRFs for *Speech-in-Quiet* vs. *Speech-in-Noise at SNR2* at the whole-brain level (p < 0.05, corrected for two tails). Lateral and medial views of the left (LH) and right (RH) hemispheres, displayed at the time-lag corresponding to the peak in the temporal profile (i.e., 250 ms). Bright yellow (positive t-values) indicates greater activation for Quiet over SNR2. Black contours indicate the ROIs borders (V1 and V2) in both hemispheres based on the Destrieux cortical atlas. **(E)** Activations obtained at the source space in visual ROIs. Boxplots display source activation for each condition, *averaged* over the ROIs (V1 + V2) and across the 0 to 600 ms time window, in the left (LH) and right (RH) hemispheres, respectively. The line through the boxplot indicates the median, × marker indicates the mean, lines indicate pairwise statistical comparisons (*p < 0.05, one-tailed).

To access the effect of SNR on neural tracking of the speech envelope, we compared the TRFs of *Speech-in-Quiet* and *Speech-in-Noise at SNR2* (the most challenging) conditions (Figure 2). The Cluster-based permutation test revealed significant differences between the TRFs for the two conditions (p < 0.025; cluster-corrected). A positive (p = 0.002, corrected) and a negative (p = 0.002, corrected) clusters were identified at time-lags interval 150 – 250 ms. Other pair of positive (p = 0.001, corrected) and negative clusters (p = 0.002, corrected) were also found at time-lags interval 290 – 410 ms (Figure 2C). Both effects extended over fronto-central and parieto-occipital electrodes. Results showed that TRF to *Speech-in-Noise at SNR2* was delayed and increased in magnitude compared to *Speech-in-Quite condition* (Figure 2A, B and C).

#### Low-level acoustic (SNR) effect at the source level

The cluster-based permutation test, performed at the whole brain level, contrasting TRFs for *Speech-in-Quiet* vs. *Speech-in-Noise at SNR2*, revealed that SNR effect was localized in both hemispheres (Figure 2D): a significant cluster was found in the left hemisphere (p = 0.008, corrected), lasting from ∼ 0 to ∼ 484 ms, and another one in the right hemisphere (p = 0.028, corrected), lasting from ∼ 141 to ∼ 312 ms (see Supplementary Figure S2). The effect was observed mostly over bilateral temporal cortex, and also included parts of the bilateral parietal cortex, insular cortex, visual cortex, and left prefrontal cortex.

#### Visual cortex ROIs

To test whether and how the visual cortex is taking part in neural tracking of speech and speech comprehension in blindfolded individuals, we performed source analysis on TRFs, using predefined ROIs in the visual cortex comprising V1 and V2.

The contrast *Speech-in-Quiet* vs. *Speech-in-Noise at SNR2* survived cluster-correction for multiple-comparisons in the left (p = 0.008, corrected) and right hemispheres (p = 0.028, corrected; Supplementary Figure S3). Extracted time-series from V1 and V2 showed a similar pattern, with the magnitude of source activation for TRF in *Speech-in-Quiet* being larger than for TRF in *Speech-in-Noise at SNR2* at multiple time points (see Supplementary Figure S3 reporting uncorrected results). Averaged activation across time points in combined ROIs (V1 + V2) was significantly larger for TRF in *Speech-in-Quiet* than for TRF in *Speech-in-Noise at SNR2* in the right hemisphere (p = 0.04, one-tailed), but not in the left hemisphere (p = 0.08, one-tailed) (Figure 2E). These results suggest the dampening of visual cortex activity in case of challenging auditory inputs.

#### High-level linguistic (Semantic) effect at the sensor level

At the two electrodes of interest (The TRFs for *Speech-in-Noise at SNR1* and *Jabberwocky-in-Noise at SNR1* significantly differed from the “null” TRF (p < 0.05, one-tailed, FDR-corrected), suggesting that the estimated TRFs indeed reflected neural tracking of the speech envelope (Supplementary Figure S1).

To access the effect of semantic information on neural tracking, we compared the TRFs of *Speech-in-Noise at SNR1* and *Jabberwocky-in-Noise at SNR1* conditions (Figure 3). Cluster-based permutation test on TRFs revealed statistically significant differences between two conditions (p < 0.025; corrected). Three pairs of positive and negative clusters were identified at time-lags intervals of 70 – 165 ms (positive: p = 0.001, corrected; negative: p = 0.01, corrected), 200 – 290 ms (positive: p = 0.001, corrected; negative: p = 0.001, corrected), and 310 – 430 ms (positive: p = 0.003, corrected; negative: p = 0.01, corrected), comprising fronto-central electrodes and parieto-occipital electrodes (Figure 3C). Results revealed that the TRFs of *Speech-in-Noise at SNR1* was higher and delayed compared to the *TRF of Jabberwocky-in-Noise at SNR1* (see Figure 3A, B and C).

**Figure 3.**
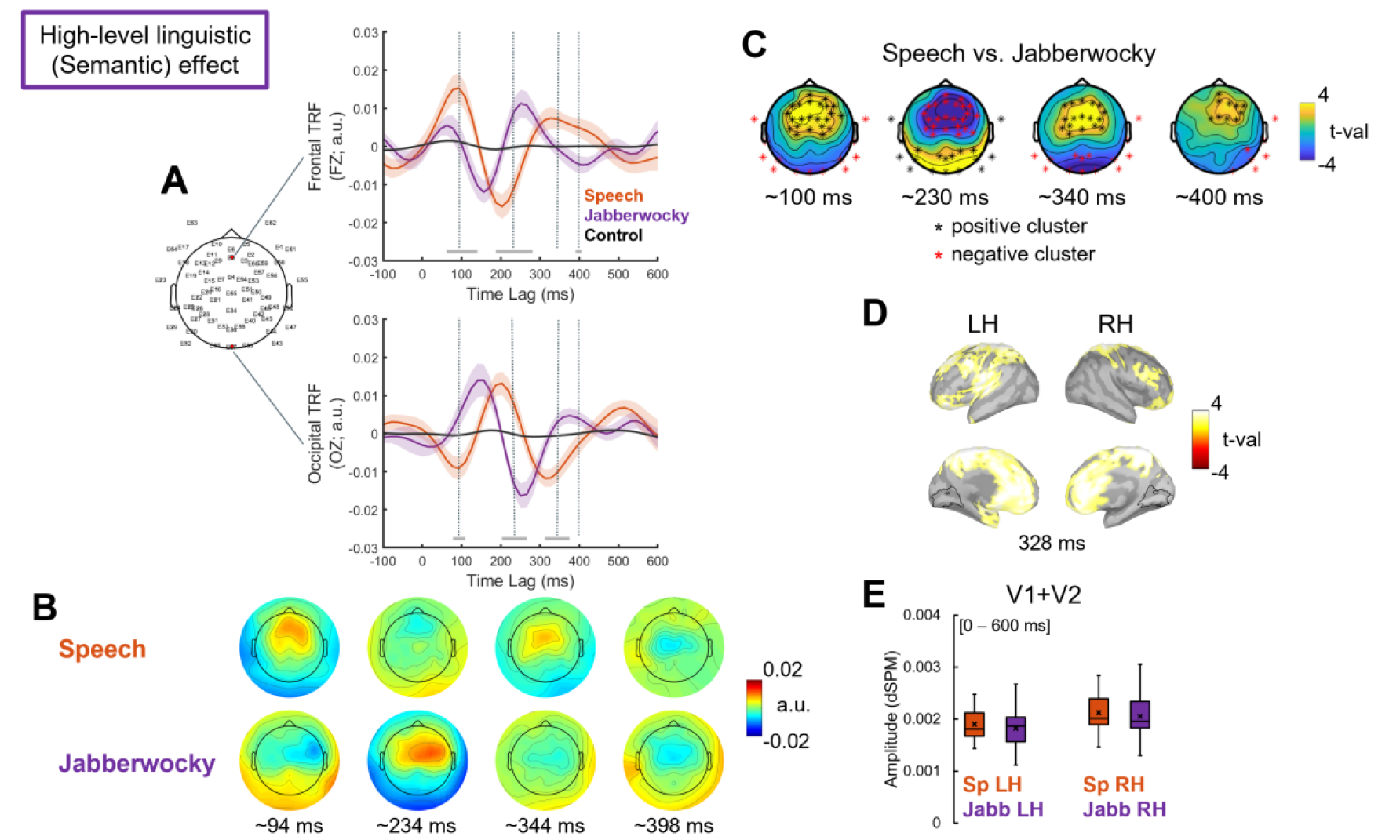
High-level (Semantic) effect. **(A)** Grand averaged temporal response functions (TRFs) for *Speech-in-Noise at SNR1* (Speech, red), for *Jabberwocky-in-Noise at SNR1* (Jabberwocky, purple), and “null” TRF (Control, black). TRFs displayed over time-lags at frontal Fz and occipital Oz electrodes, marked with red on the electrode layout. Shaded areas represent the standard error of the mean (SE) across participants. Grey horizontal bars above the x-axis indicate time-lags at which TRFs for *Speech-in-Noise at SNR1* and *Jabberwocky-in-Noise at SNR1* differed significantly (running paired two-tailed t-tests, p < 0.05, FDR-corrected). Grey dotted vertical lines indicate time-lags with the maximal difference between TRFs for *Speech-in-Noise at SNR1* and *Jabberwocky-in-Noise at SNR1*. **(B)** Topographic representations of TRFs, displayed at time-lags marked by grey dotted vertical lines on **A. (C)** The results of the cluster-based permutation test contrasting TRFs for *Speech-in-Noise at SNR1* and *Jabberwocky-in-Noise at SNR1*, displayed around time-lags marked by grey dotted vertical lines on **A**. Significant (p < 0.05, cluster-corrected for two tails, meaning p < 0.025 each tail) positive and negative clusters comprised the electrodes marked in black and in red asterisks, respectively. **(D)** Differences at the source level, contrasting TRFs for *Speech-in-Noise at SNR1* and *Jabberwocky-in-Noise at SNR1* at the whole brain level (p < 0.05, cluster-corrected for two tails). Lateral and medial views of the left (LH) and right (RH) hemispheres, displayed at the time-lag corresponding to the peaks in the temporal profile. Bright yellow (positive t-values)/dark red (negative t-values) colors indicate greater activation for Speech/Jabberwocky, respectively. Black contours indicate the ROIs borders (union of V1 and V2) in both hemispheres based on the Destrieux cortical atlas. **(E)** Activations obtained at the source space in visual ROIs. Boxplots display source activation for each condition, *averaged* over the ROIs (V1 + V2) and across the 0 to 600 ms time window in the left (LH) and right (RH) hemispheres, respectively. The line through the boxplot indicates the median, × marker indicates the mean.

#### High-level linguistic (Semantic) effect at the source level

Cluster-based permutation test, contrasting TRFs for *Speech-in-Noise at SNR1* and *Jabberwocky-in-Noise at SNR1* at the whole-brain level, revealed two clusters in both hemispheres: one in the left hemisphere (p = 0.002, corrected), extending over all time points, and one in the right hemisphere (p = 0.006, corrected), lasting from ∼0 to ∼531 ms (Supplementary Figure S2), with maximum activation ∼ 330 ms (Figure 3D). The effect extended primarily over the left auditory cortex and a large portion of the bilateral fronto-parietal network at earlier time points and extended to the anterior temporal lobe (ATL) at later time points (Supplementary Figure S2).

#### Visual cortex ROIs

In the visual ROI the Semantic effect did not survive cluster-correction for multiple-comparisons (p > 0.05, corrected for two tails), and extracted time-series from ROIs did not differ between source TRFs for *Speech-in-Noise at SNR1* and *Jabberwocky-in-Noise at SNR1* (p > 0.05) (Supplementary Figure S3 and Figure 2E).

## Discussion

We used a hierarchical model to investigate entrainment to continuous speech envelope in blindfolded individuals, assessing (1) the effects of low-level acoustic and high-level linguistic information on neural tracking and (2) testing how these effects are distributed at the source level, with the focus on the visual cortex. To address the role of low-level acoustic, we compared the entrainment to target stories presented in quiet or multi-talker babble noise. Results revealed that TRF was delayed and higher in magnitude at latencies between 100 and 300 ms when SNR decreased. This finding suggests that neural tracking requires greater resources in case of concurrent masking noise. Next, we also addressed the role of high (semantic) level of speech processing on neural entrainment by comparing TRFs to meaningful and meaningless stories. Results indicated delayed and higher TRFs when semantic information is present. Source modeling suggested that entrainment to continuous speech in noise engaged a spread activation beyond the auditory cortex, including linguistic and attentional networks. Finally, in the absence of retinal input, we found evidence that the visual cortex entrained to the speech envelope. However, the magnitude of such entrainment was degraded with concurrent background noise, suggesting a suppressing mechanism helping to focus auditory attention in challenging listening conditions.

### Effects of low-level acoustic (SNR) processing on neural tracking of speech envelope

We demonstrated that speech envelope tracking in noise, compared to quiet, was characterized by larger amplitude and delayed latency of the TRF responses and by the reversed polarity of the TRFs topography distributions over fronto-central parieto-occipital electrodes (Figure 2A, B).

The TRF time-courses were consistent with previous studies reporting amplitudes and latencies being affected by concurrent noise (Brodbeck, Jiao, Hong, & Simon, 2020; Ding & Simon, 2013; Fiedler, Wöstmann, Herbst, & Obleser, 2019; Gustafson, Billings, Hornsby, & Key, 2019; Zendel, West, Belleville, & Peretz, 2019) as well as enhanced N1 and N2 amplitudes in noise compared to quiet (Papesh, Billings, & Baltzell, 2015).

Increased frontal negativity around 100 ms (N1) is associated with attention-dependent processes in response to auditory changes (Hansen & Hillyard, 1980; Näätänen, 1982). The enhanced envelope tracking observed here for the N1-like response to speech in noise compared to quiet may reflect the use of more resources for the encoding of acoustic variations at earlier stages of speech processing when intelligibility gets degraded by noise (Alain, Quan, McDonald, & Van Roon, 2009; Näätänen & Picton, 1987; Parbery-Clark, Marmel, Bair, & Kraus, 2011).

Additional differences were observed around the second negative peak, corresponding to the N2 component. The TRF peak around this component was smaller and delayed for speech in noise compared to speech in quiet. Delayed N2 response is associated with attentive speech processing in challenging acoustic conditions (Balkenhol, Wallhäusser-Franke, Rotter, & Servais, 2020; Billings, Tremblay, Stecker, & Tolin, 2009; Finke, Büchner, Ruigendijk, Meyer, & Sandmann, 2016). Again, differences in this time range (between 100 and 300 ms after stimulus onset) possibly reflect changes in the degree of attention required to encode incoming stimuli effectively. Particularly, delayed TRF peak response may reflect participants’ effort in keeping track of meaningful information over time in the degraded signal. Compensatory mechanisms may be involved in segregating speech from noise. Previous evidence reported stronger envelope tracking of attended speech with increased background noise in hearing-impaired and elderly individuals compared to hearing younger adults (Brodbeck, Presacco, Anderson, & Simon, 2018; Decruy, Vanthornhout, & Francart, 2020; Presacco, Simon, & Anderson, 2016). Both internal (hearing loss) and external (background noise) factors can produce acoustic distortion, which may result in increased listening effort (Van Engen & Peelle, 2014) and enhanced envelope tracking.

There is a debate whether envelope tracking is enhanced (Ding et al., 2014; Ding & Simon, 2013; Fuglsang, Dau, & Hjortkjær, 2017; Presacco et al., 2016) or reduced (Desai et al., 2021; Ding & Simon, 2013; Kurthen et al., 2021; Vanthornhout, Decruy, Wouters, Simon, & Francart, 2018; L. Wang, Wu, & Chen, 2020) with decreasing SNR. Our behavioral results showed that comprehension scores and intelligibility rates were directly proportional to SNR levels. Our results on TRFs also add to the findings that envelope tracking increases with noise and when listening becomes more challenging.

### Effects of high-level linguistic (Semantic) processing on neural tracking of speech envelope

Topographical distributions of the TRFs suggest the involvement of distinct neural generators when semantic content is present or absent (Figure 3B). Moreover, the temporal dynamics of TRFs for meaningful story was characterized by a more prominent P1 peak and generally delayed P1-N1-P2-N2 complex, as compared to meaningless story (Figure 3A).

At a relatively early processing stage (around 100 ms), we observed stronger neural tracking of the speech envelope for meaningful story than for meaningless story over fronto-central electrodes (Figure 3A, B). This finding could seem surprising since auditory P1 is often associated with pre-attentive processes such as onset detection and sensory gating (Huotilainen et al., 1998; Miller, Graham, & Schafer, 2021; Thoma et al., 2003; Waldo et al., 1992). Predictive models of speech processing provide a plausible explanation for this result. Semantic content generates expectations about upcoming stimuli and limits the degree of uncertainty about what was heard (Poeppel, Idsardi, & van Wassenhove, 2008), affecting early auditory encoding (Broderick, Anderson, & Lalor, 2019) and neural tracking of the speech envelope (Di Liberto et al., 2018; Kaufeld et al., 2020). Meaningful information may provide regularities in meaningful story, making it more predictable than meaningless story.

Moreover, it is possible that envelope tracking of meaningless story may not be affected by the background noise as much as meaningful story due to the difference in the degree of informational masking. It is possible that meaningless story could “pop-out” from the background multi-talker babble noise due to lower informational masking compared to meaningful story. Under the linguistic similarity hypothesis (Van Engen & Bradlow, 2007), informational masking is more efficient when background babble noise has more linguistic similarity with the target speech stream (e.g., same spoken language, known accent) compared to a different or unknown language, accent and semantically anomalous speech (Brouwer, Van Engen, Calandruccio, & Bradlow, 2012; Brungart, 2001; Calandruccio, Van Engen, Dhar, & Bradlow, 2010; Cooke, Garcia Lecumberri, & Barker, 2008; Garcia Lecumberri & Cooke, 2006; Van Engen, 2010; Van Engen & Bradlow, 2007). Therefore, it could have been easier for participants to segregate from the background noise meaningless story than meaningful story.

### Two distributed networks are engaged in envelope tracking of continuous speech

Source analysis of TRFs highlighted temporal and fronto-parietal regions traditionally involved in speech and language comprehension (Hertrich, Dietrich, & Ackermann, 2020). Key regions for low-level acoustic effect tested here involved the bilateral temporal cortex, parts of the parietal, insular, and visual cortices bilaterally, and the left prefrontal cortex (Figure 2D). Naturalistic speech stimuli are complex and resemble everyday listening conditions, thus leading to extended activations and involvement of higher-order cortical regions (Alexandrou, Saarinen, Kujala, & Salmelin, 2020; Hamilton & Huth, 2020). For example, narrative speech involves widely distributed bilateral neural activity that tracks hierarchically organized speech representations at multiple cortical sites and temporal windows (de Heer, Huth, Griffiths, Gallant, & Theunissen, 2017; Di Liberto et al., 2015; Huth, de Heer, Griffiths, Theunissen, & Gallant, 2016; Lerner, Honey, Silbert, & Hasson, 2011; Poeppel, 2003; Puschmann, Regev, Baillet, & Zatorre, 2021). Neuroimaging studies reported distributed cortical activations beyond the auditory cortex (comprising higher-order associative brain structures and attentional networks) during effortful listening (see Alain, Du, Bernstein, Barten, & Banai, 2018 for a meta-analysis).

Higher-level linguistic processing was assessed by contrasting meaningful and meaningless stories (Speech vs. Jabberwocky) and resulted in higher activation for meaningful story, mainly involving the left auditory cortex, a large portion of bilateral fronto-parietal network, and the left anterior temporal lobe later in time (Figure 3D). Overall source modeling results of TRFs indicate that low-level acoustic effects mainly involved a bilateral temporo-parietal network, while higher-level linguistic effects primarily involved a left dominant fronto-temporal network. These results support the notion that successful speech comprehension requires multiple extended networks beyond the temporal lobe to process the acoustic signal at multiple and parallel hierarchical levels (Davis & Johnsrude, 2003, 2007; de Heer et al., 2017; Hickok & Poeppel, 2007; Peelle, 2012; Peelle, Johnsrude, & Davis, 2010)

### Early visual cortex’s entrainment to speech envelope in blindfolded individuals is reduced by background noise

We performed source analysis on the TRFs from preselected visual ROIs (V1 and V2) to assess whether the visual cortex contributes to neural envelope tracking in blindfolded individuals. While source estimates of EEG activity should be taken with caution, results suggested early visual cortex’s involvement in envelope tracking, especially for low-level acoustic speech processing (Figure 3D, E).

A recent fMRI study showed that the visual cortex of blindfolded individuals displayed some degree of synchrony to audio tracks from movies and narratives, suggesting that auditory information can reach the visual cortices (Loiotile, Cusack, & Bedny, 2019). Overall, numerous fMRI findings supported the notion that the visual cortex is functionally engaged in processing non-visual stimuli in sighted individuals (Facchini & Aglioti, 2003; Merabet et al., 2008; Poirier et al., 2006; Qin & Yu, 2013; Ricciardi et al., 2011; Sathian, 2005; Seydell-Greenwald et al., 2021; Vetter et al., 2014; Zangaladze, Epstein, Grafton, & Sathian, 1999).

Interestingly, we observed a decrease in total signal magnitude for speech in noise compared to speech in quiet. This difference emerged in particular for the right visual cortex (although a trend also existed in the left hemisphere; Figure 2E). Hemispheric asymmetry is not surprising, as previous evidence already showed the right hemisphere dominance for several aspects of natural speech processing, especially for tracking of slow temporal modulations within the delta-theta range (Alexandrou, Saarinen, Mäkelä, Kujala, & Salmelin, 2017; Poeppel, 2003). More importantly, this finding aligns with the evidence that the early visual cortex is sensitive to acoustic SNR effects (Bishop & Miller, 2009).

These results seem to suggest that the activity of the visual cortex could be modulated during continuous speech tracking. However, its activity gets suppressed if the attentional network becomes more engaged in tracking relevant auditory information in a challenging listening environment. Human neuroimaging studies reported cross-modal deactivation of the visual cortex by auditory stimuli during active listening or passive stimulation (with the instructions to concentrate on the stimuli) and suggested that such suppression can be top-down modulated by attention as task demands increase (e.g., Hairston et al., 2008; Johnson & Zatorre, 2006; Laurienti et al., 2002). Several other studies found suppression effects of sound on visual perception. Such cross-modal suppression has been suggested to reduce the magnitude of the percept of a weaker or less relevant modality input considered as a perceptual noise (Hidaka & Ide, 2015).

Overall, our results align with recent evidence reporting that the visual cortex can contribute to auditory information processing in sighted individuals (Brang et al., 2022; Martinelli et al., 2020; Seydell-Greenwald et al., 2021; Vetter et al., 2014). Here, we observed that the visual cortex is more engaged in processing when speech signal is intelligible and clear (i.e., presented in quiet). Differences in mapping speech envelope in the visual cortex for low-level acoustic representations exist and might reflect cross-modal visual cortex suppression. Such suppression could be top-down modulated and attributed to auditory attention (Cate et al., 2009), which plays an essential role in segregating relevant speech information in challenging listening conditions and when congruent visual input is unavailable.

It could be argued that mental imagery mechanisms may drive the visual cortex’s response to speech. Previous studies observed an overlap in neural representations in the occipital areas between perception and visual imagery, stemming from common top-down influences (see Dijkstra, Bosch, & Gerven, 2019 for a review). However, V1 has been shown to encode auditory information regardless of imageability (Martinelli et al., 2020; Seydell-Greenwald et al., 2021; Vetter et al., 2020, 2014). Thus, the role of the early visual cortex in auditory processing may not be merely ascribed to an imagery effect. If that was the case, when contrasting *Speech-in-Noise* and *Jabberwocky-in-Noise*, we could have observed higher visual cortex’s responses in meaningful condition compared to meaningless one, since only the former contained visually imaginable information. However, no significant difference in the visual cortex’s entrainment to speech envelope was found between these conditions.

### Limitations and future research perspectives

It is important to acknowledge the challenges of EEG-based source modeling, as spatial resolution of EEG is generally known to be relatively poor, making it difficult to identify exact brain sources that generate the neuronal activity measured on the scalp. EEG-based source modeling majorly suffers from an ill-posed inverse problem and can also result in misleading activity patterns due to, for instance, low SNR, unrealistic head models, invalid constraints, and so on. More accurate EEG source localization requires digitized electrode positions and individual anatomical scans of participants, which can diminish source estimation uncertainty (Shirazi and Huang, 2019; Michel and Brunet, 2019; Zorzos et al., 2021) but were not available in our study. Therefore, EEG source estimates should be interpreted with caution. However, it is worth noting that we used a validated pipeline for source modeling estimation (Stropahl et al., 2018; Bottari et al., 2020). Moreover, the same source modeling was performed across different conditions; thus, similar errors should be attributed to activations for each condition. While the exact location of the activity cannot be ensured with the present data, our results suggested that the activity of posterior cortices was modulated only by low-level and not high-level speech processing.

A further limitation pertains the input data we used for the encoding. We modeled neural tracking of the speech signal based on a single feature: the speech envelope comprising specific bandwidth frequencies (2-8 Hz). The envelope represents slow-variate temporal modulations of the speech signal. It contains multiple acoustic and linguistic cues important for continuous speech segmentation into smaller units, and therefore it has been hypothesized to be crucial for speech comprehension (Luo & Poeppel, 2007; Shannon, Zeng, Kamath, Wygonski, & Ekelid, 1995; Zoefel, 2018). However, it has also been argued that focusing on the envelope alone might not get the complete picture of the neural mechanism underlying speech comprehension (Obleser, Herrmann, & Henry, 2012). Recent studies reported that the inclusion of multiple speech features, such as spectrogram, phonemes, and phonetic features in the model sometimes result in a better model performance represented by a more robust neural tracking response (e.g., Brodbeck, Hong, & Simon, 2018; Di Liberto et al., 2015, 2018; Lesenfants, Vanthornhout, Verschueren, Decruy, & Francart, 2019). Future research may include multiple speech features to build a multivariate model to assess neural speech tracking in the brain and how the visual cortex maps speech information when visual input is absent.

## Conclusion

Overall, our results indicate low-level acoustic and high-level linguistic processes affecting envelope tracking of continuous speech. Envelope tracking may play a role in supporting active listening in challenging conditions and is enhanced when SNR decreases, and when segregation of target speech from the background noise becomes more difficult (i.e., due to linguistic similarity). Tracking speech signal embedded in noise requires spread networks of activation, including linguistic and attentional regions beyond the auditory cortex. In the absence of retinal input, the visual cortex might entrain to the speech envelope, however, the functional role of such activity remains to be ascertained. The magnitude of entrainment is degraded by concurrent noise, suggesting a suppressing mechanism aimed at focusing resources within the auditory attention network in case of challenging listening conditions. Conversely, no clear impact of semantic content was found in the visual cortex, suggesting that the magnitude of such entrainment is generally affected by low-level speech features.

## Acknowledgements

The authors thank Chiara Battaglini who helped with stimuli recordings and preparation. We also thank Chiara Maccioni for lending her voice for stimuli recordings. Davide Bottari is funded by PRIN 2017 research grant. Prot. 20177894ZH.

## Author contributions

Conceptualization, D.B., E.B., B.M., S.D.; Methodology, D.B., E.B., A.M.; Formal Analysis, E.B., B.M., D.B.; Investigation, E.B, A.M.; Data Curation – E.B.; Writing – Original Draft, E.B., M.B., A.F., D.B.; Writing – Review and Editing, E.B., D.B., M.B., A.F., B.M., S.D., E.R., A.M.; Visualization, E.B., D.B.; Resources & Funding, D.B.

