## Supplementary Material for "Early visual cortex tracks speech envelope in the absence of visual input"

### Questionnaire

Readers can contact the corresponding author to request access to this information.

### Choice of SNR level

The range of SNR levels was based on the results of our pilot experiments, with the idea of being capable to considerably affect participant’s intelligibility rather than severely disrupting it. To select SNR levels, we behaviorally tested six other participants who did not take part in the main study. They listened short (~ 1 min) speech fragments embedded in noise at a range of fixed SNR levels ({-3.52 dB, -1.74 dB, 0 dB, +1.74 dB, + 3,52 dB}), with randomly drawn parts of the babble noise on the background to prevent participants' adaptation to a particular babble. As a result, we identified two SNR levels: +3.52 dB (SNR1, as an easier level for participants to comprehend) and +1.74 dB (SNR2, as a harder level for participants to comprehend).

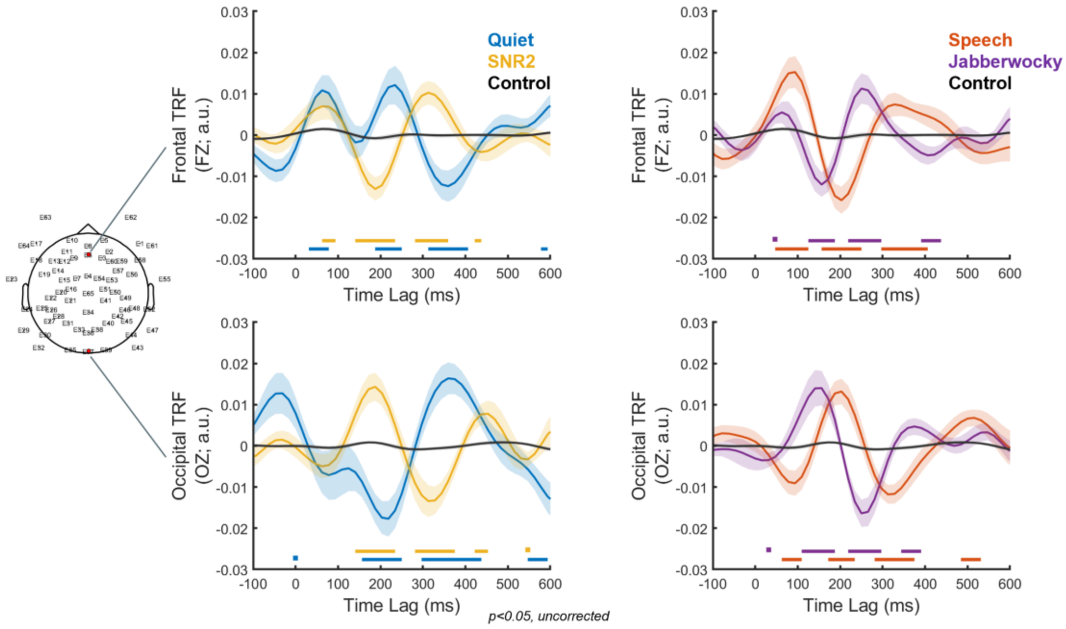


**Supplementary Figure S1**: **TRF models’ performance.** Grand averaged temporal response functions (TRFs) for four experimental conditions: *Speech-in-Quiet* (Quiet, blue), *Speech-in-Noise at SNR2* (SNR2, yellow), *Speech-in-Noise at SNR1* (Speech, red), *Jabberwocky-in-Noise at SNR1* (Jabberwocky, purple); and "null distributed" TRF model (Control, black). TRFs displayed over time-lags at frontal Fz and occipital Oz electrodes, marked with red on the electrode layout. Shaded areas represent the standard error of the mean (SE) across participants. Colored horizontal bars above the x-axis indicate time-lags at which TRFs of experimental conditions differed from the "null TRF" ($p < 0.05$, uncorrected).

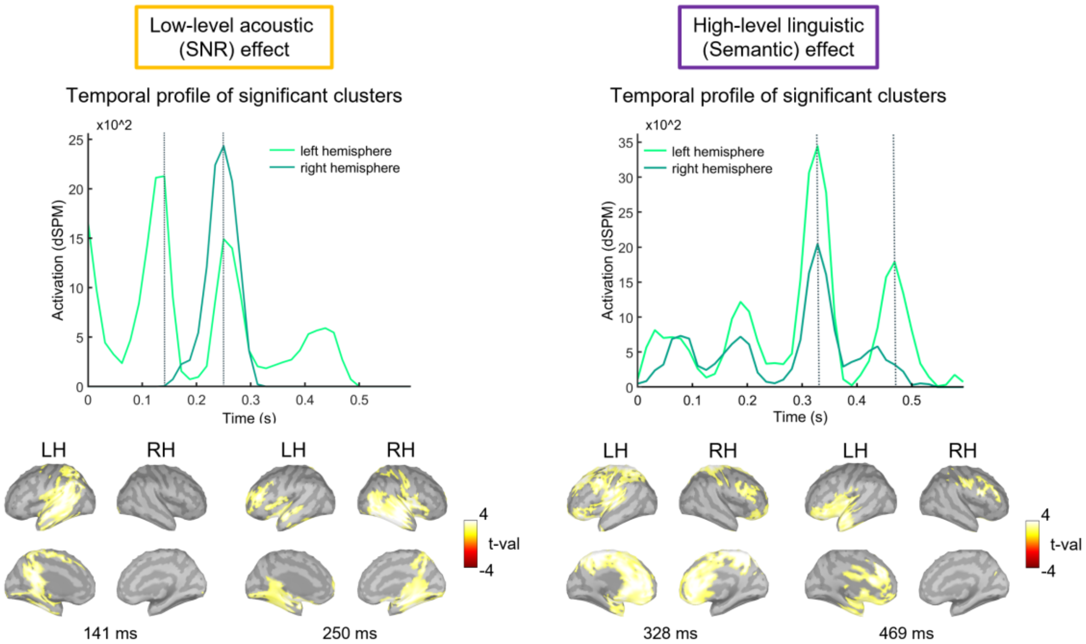


**Supplementary Figure S2**: **Source analysis, whole-brain level.** Grand averaged TRF source-localization results, corrected for multiple comparisons using a cluster-based permutation test across all electrodes and time-lags from 0 to 600 ms. Top panel: Temporal extent profile for each of the identified significant clusters in the left (light green) and right (dark green) hemispheres. Grey vertical lines mark time-lags corresponding to visually salient peaks in the temporal profile, in the left and right hemisphere, respectively. Bottom panel: Significant differences between TRFs at the source space ($p < 0.05$, cluster-corrected); lateral and medial views of the left (LH) and right (RH) hemispheres, displayed at the time-lags marked on the Top panel. Bright yellow/dark red colors indicate greater activity for Quiet/SNR2 and Speech/Jabberwocky, respectively.

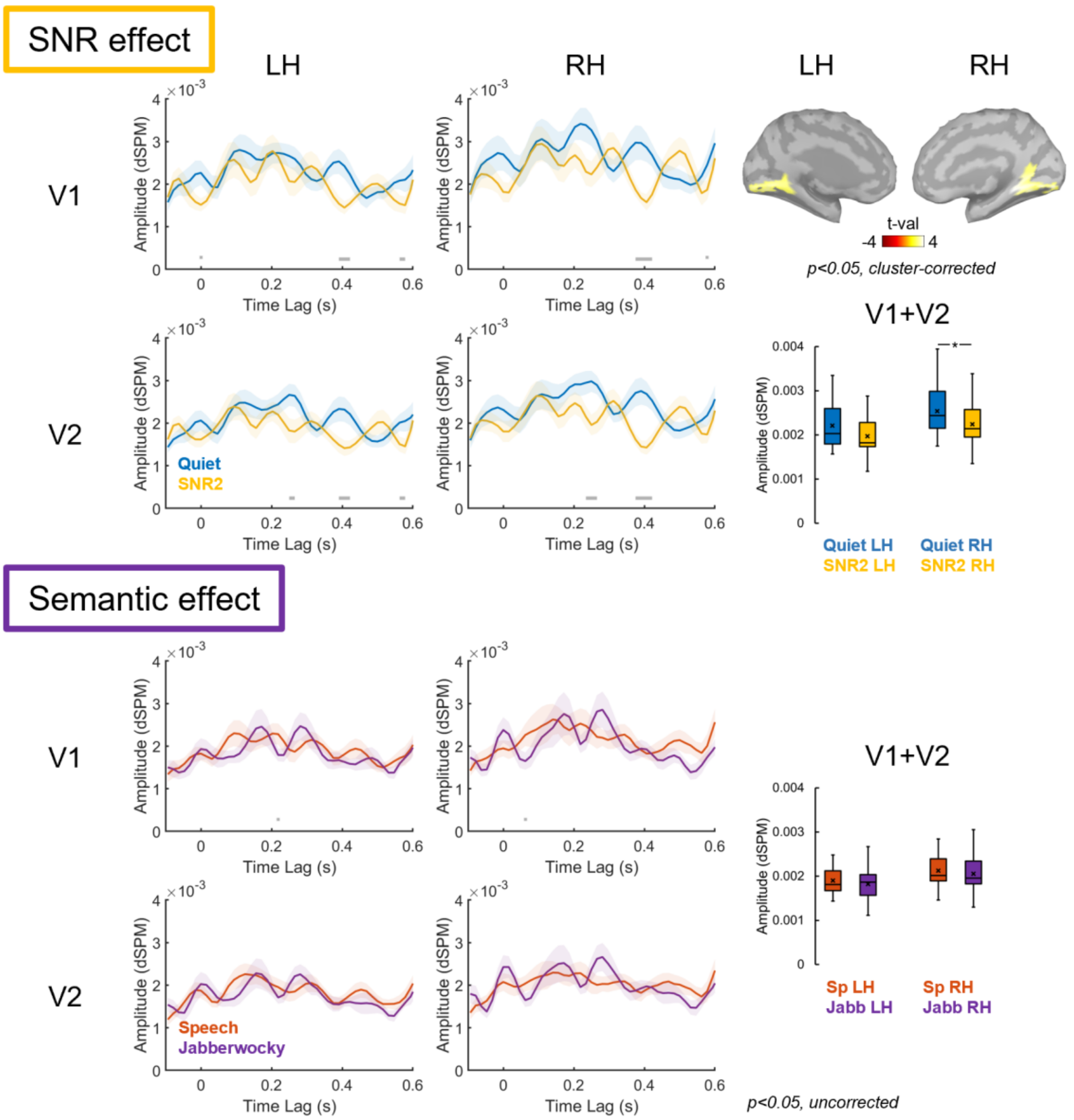


**Supplementary Figure S3: Source analysis, visual ROIs.** Grand averaged time series of the activation extracted for visual ROIs (V1 and V2), for the left (LH) and right (RH) hemispheres. Activation is displayed in unitless and absolute values, as provided by the normalization within the dSPM algorithm. Shaded areas indicate the standard error of the mean (SE). Grey horizontal bars above the x-axis indicate time-lags at which two conditions significantly differed ($p < 0.05$, uncorrected). Statistical map projected on the brain surface to highlight differences between conditions (Quiet vs. SNR2; $p < 0.05$, cluster-corrected) in ROIs, identified at the whole-brain level and displayed for the left (LH) and right (RH) hemispheres at ~ 250 ms (corresponding to the peaks in the temporal profile); bright yellow/dark red colors indicate greater activity for Quiet/SNR2, respectively. Boxplots display source activation for each condition, *averaged* over the ROIs (V1 + V2) and across all relevant time-points (from 0 to 600 ms regarding speech envelope onset), in the left and right hemispheres, respectively. The line through the box indicates median; × marker indicates the mean; lines indicate pairwise statistical comparisons (*p < 0.05).

### Effect of Speech Rate, Intensity and Modulation Depth

Neural tracking could be affected by speech rate (Müller et al., 2019) and intensity of the stimuli (Drennan & Lalor, 2019). Thus, it could be argued that our results could also be driven by such differences. However, the target speaker was the same in all conditions, and RMS amplitude of the stimuli was normalized to a constant value, allowing to control for both speech rate and intensity.

Speech presented in quiet varies in amplitude modulation depth because of silent gaps, whereas speech presented in noise has homogenous amplitude. If amplitude modulation depth would have an effect, we should have observed clear between-condition differences in the early peak (P1), that has been suggested to reflect early, pre-perceptual sound processing (Ceponienė et al., 2005), which was not the case.

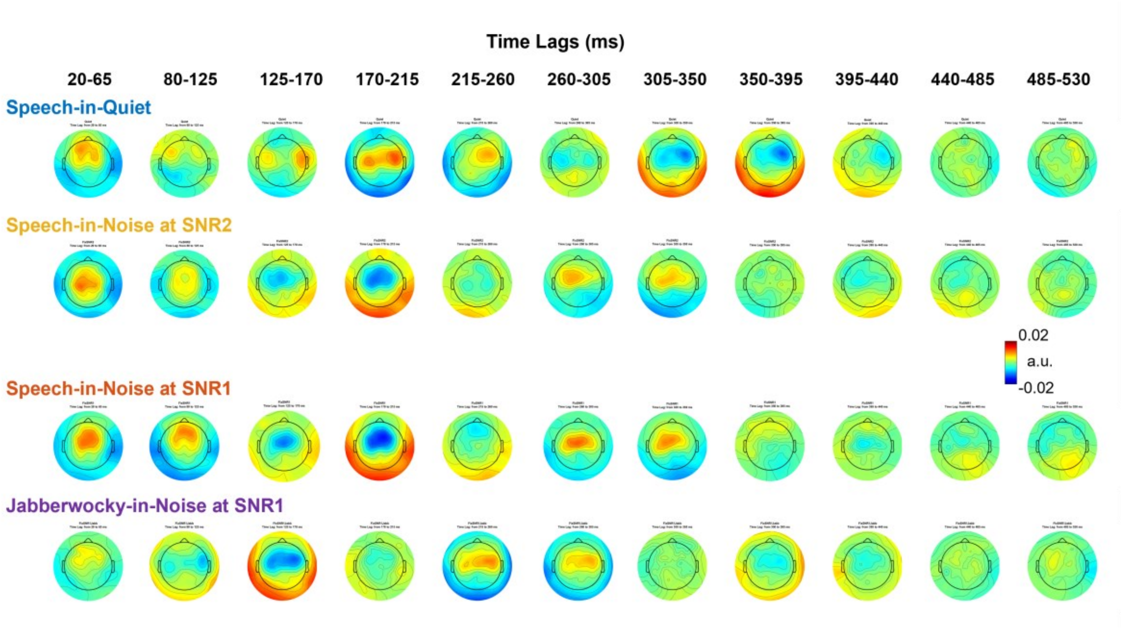


**Supplementary Figure S4: Dynamics of the TRFs in time.** Topographic representations of grand averaged temporal response functions (TRFs), displayed over multiple time-lags for each experimental condition.
